# Songs in the understory and colors in the canopy: habitat structure leads to different avian communication strategies in a tropical montane forest

**DOI:** 10.1101/393595

**Authors:** Oscar Laverde-R, Carlos Daniel Cadena

## Abstract

Birds inhabit a variety of habitats and they communicate using primarily visual and acoustic signals; two central hypotheses have been postulated to study the evolution of such a signals. The sensory drive hypothesis posits that variation in the physical properties of habitats leads to variation in natural selection pressures by affecting the ease with which different types of signals are perceived. Assuming that resources are limited for animals, the transfer hypothesis predicts a negative relationship between the investments in different types of signals. We evaluated these two hypotheses in a tropical montane forest bird assemblage. We also postulate a possible interaction between these two hypotheses: we predicted that the negative relationships between signals should be observed only when jointly considering birds from different environments (e.g. understory and canopy) due to the expected differences in communication strategies between habitats. The sensory drive hypothesis was supported by the differences we found between strata in vocal output, patch contrast to background and color conspicuousness, but not for the variables associated to song elaboration and hue disparity. We found support for the transfer hypothesis: birds with colors contrasting less against the background sing more frequently and birds with lower diversity of colors produce longer songs, understory birds showed also a negative relationship between signals, but only when accounting for phylogeny. We found partial support for the interaction between the sensory drive and the transfer hypotheses: hue disparity and vocal output were negatively related only when analyzing together birds from the canopy and the understory, but not when analyzing them separately. We conclude that the study of the evolution of communication signals needs to consider more than one channel and the functional interactions between them. The results of the interaction of optimal signaling strategies in two communication channels in the local habitats where animals signaling, are the patterns of colors and songs we revealed in a tropical montane forest bird assemblage.

## Introduction

Birds communicate using primarily visual and acoustic signals, and the evolution of these signals likely involves a compromise between sexual selection for conspicuousness and natural selection for crypsis (Endler 1992). Assuming that producing communication signals (i.e. plumage colors, songs) is costly and that resources devoted to producing such signals are limited, one expects that animals should invest predominantly in one type of signal (Shutler, 2011). Accordingly, the trade-off or transfer hypothesis predicts a negative relationship between the investment in different types of signals: birds with complex songs are expected to exhibit dull plumages, whereas birds with colorful plumages are expected to have simple songs (Darwin 1871; Huxley 1938; Gilliard 1956; Shutler and Weatherhead 1990; Kusmierski et al. 1997; Ornelas et al. 2009; Shutler 2011). In addition, the direction in which potential trade-offs between plumage and song elaboration are resolved is likely to be affected by spatially variable biotic and abiotic factors. For example, if high predation in a given environment selects for reduced conspicuousness in plumage, then song elaboration may replace plumage characteristics as a main target of sexual selection, resulting in increased elaboration of vocal signals (Badyaev et al. 2002).

The sensory drive hypothesis posits that variation in the physical properties of the habitat in which animals communicate leads to variation in natural selection pressures by affecting the ease with which different types of signals are perceived (Endler 1992). The canopy and understory of tropical forests, for example, are highly contrasting habitats in terms of light availability (Gomez and Théry 2004, 2007) and acoustic properties affecting sound transmission (Ryan and Brenowitz 1985; Nemeth et al. 2001; Slabbekoorn and Smith 2002; Seddon 2005). Therefore, one may hypothesize that birds occupying different forest strata are likely to be differentially selected with regard to colors and songs (Nemeth et al. 2001; Gomez and Théry 2004, 2007; Seddon 2005).

The forest understory is a dark environment in which ultraviolet (UV) light is especially scarce, and in which the visual background is mostly brown due to tree trunks and leaf litter (Endler 1993). In contrast, light is abundant and rich in UV wavelengths in the forest canopy, where the background is largely green due to leaves. Work on an avian assemblage inhabiting a Neotropical lowland forest suggests that the plumage patterns of birds vary between understory and canopy species in ways consistent with adaptation to contrasting light environments (Gomez and Théry 2007). For example, UV patches and colors rich in short wavelengths (i.e. blue and violet) are relatively frequent in the plumage of canopy birds but are nearly absent in the understory. Understory birds are usually monochromatic and often exhibit brown dorsal plumage showing little contrast to their background, whereas conspicuous yellow and orange are more common colors in ventral plumages. In contrast, the most common colors in the canopy are green, black and brown in males, and brown and green in females. The contrast in brightness between dorsal plumage patches and the background is greater in canopy species than in understory species. In sum, under the avian visual system, canopy birds are more conspicuous relative to their background than understory birds in a lowland tropical forest (Gomez and Théry 2007). Greater conspicuousness in the canopy may originate from less intense selection by predators for crypsis or from the greater efficiency of UV signals in this environment, which allows for such signals being used for communication with conspecifics while being undetected by some predators (i.e., a private communication channel; Gomez and Théry 2007). Whether this applies to other forest sites remains to be seen.

As with colors, songs and calls are thought to be shaped by habitat structure. The acoustic adaptation hypothesis postulates that structural properties of habitats influence signal evolution through their effects on signal transmission (Jønsson et al., 2012; Kirschel et al., 2009; Morton, 1975; Nemeth et al., 2001; Ryan & Brenowitz, 1985; Seddon, Merrill, & Tobias, 2008; Slabbekoorn & Smith, 2002; Tobias et al., 2010). Dense habitats are expected to select for lower-pitched vocalizations with narrower frequency bands and for songs with longer notes and inter-notes relative to open habitats (Morton 1975; Ryan & Brenowitz 1985). In forest environments the understory is more densely vegetated than the canopy, while stems and branches are thinner and leaves smaller in the canopy; this structural variation in the environment results in height-specific patterns of sound degradation affecting the design of vocal signals (Nemeth et al. 2001). Specifically, birds living in the understory are expected to use low-frequency songs to minimize the effects of frequency attenuation and to emit long, well-spaced notes to avoid distortion by reverberation (Nemeth et al. 2001). But some narrow frequency bandwidth vocalizations reverberations may be beneficial leading to longer and louder notes after signal transmission (Slabbekoorn et al. 2002). In the canopy, the effects of frequency attenuation and reverberation are less strong; thus, songs of canopy species are expected to be higher pitched, with shorter notes and less time between notes relative to songs of understory species (Nemeth et al. 2001).

Many comparative studies have tested and supported the sensory drive hypothesis, but these studies have largely focused only on single communication channels, emphasizing the study of either colors (Mcnaught and Owens 2002; Doucet et al. 2007; Ornelas et al. 2009; Price and Whalen 2009; Cardoso and Mota 2010) or songs (Kirschel, et al., 2009; Smith & Bernatchez, 2008; Kirschel, et al., 2009; Seddon & Tobias, 2010; Seddon, 2005; Podos, 2001; Nowicki et al., 1998; Price & Lanyon, 2002; Price, 2004). However, if there are tradeoffs in the investment between visual and acoustic signals, then the direction of evolution will depend on the influence of the environment (i.e. sensory drive) on how such tradeoffs are resolved. For example, a recent study on the evolution of communication signals in New World Warblers (Parulidae) showed that sensory drive influenced the transfer of investment between traits in different sensory modalities (Laverde et al., Chapter 3). In that study, negative relationships between measures of song and plumage elaboration were only apparent when considering the influence of habitat: plumage contrast to the background and chromatic diversity were negatively related to song syllable variety only when vegetation structure was included as a covariate in analyses (e.g., birds with a greater variety of song syllables and less colorful plumage live in closed or darker habitats). Thus, negative relationships between investment in different types of signals do not necessarily exist because of internal physiological or energetic restrictions, but may arise as a consequence of selection for optimal signaling strategies in the local habitat of species (Laverde et al., Chapter 3).

A tropical montane forest is an appropriate scenario for studying the variation in visual and acoustic signals of birds in relation to habitat structure. At a single locality, one may find a large number of bird species exhibiting a wide variety of colors and songs, and differing in microhabitat use within the forest (Stiles and Rosselli 1998). In addition, the gradient of physical properties from the canopy to the understory within a montane forest offers a good framework for testing the sensory drive and the transfer hypotheses while exploring possible interactions between them. Here, we studied an assemblage of tropical montane birds to examine whether mechanisms underlying negative relationships between signals (i.e. transfer hypothesis) may be understood as the result of interactions between suitability of different types of signals for communication given habitat features (i.e. sensory drive hypothesis; Seehausen et al. 2008; Tobias and Seddon 2009; Laverde et al., Chapter 3).

First, we tested the sensory drive hypothesis (i.e. the effect of contrasting habitats: understory vs. canopy) using variables related to song elaboration and vocal output and variables related to color diversity and conspicuousness. The hypothesis that signal evolution is driven by sensory drive predicts that in the understory, where light availability is low and visual signals are easily obstructed by vegetation, birds would rely more in acoustic than in visual signals; this should be manifested in greater vocal output (i.e. in singing more frequently) or in greater song elaboration relative to canopy species (Fig. 1a). By contrast, visual signals (i.e. colorful or contrasting plumages) are predicted to be favored in canopy birds due to greater light availability and less obstacles. However, because singing in colorful species in a more open habitat like the canopy would further increase detectability by predators (Marler 1955; Gotmark and Post 1996), one expects vocal output to be lower or songs to be less elaborate in canopy species than in understory species (Fig. 1b). Second, we tested the transfer hypothesis between songs and colors, which predicts negative relationships between song elaboration or vocal output and variables indicating plumage conspicuousness regardless of habitat (i.e. negative relationships should exist among species from the same forest stratum; Fig. 1c and 1d). Finally, we tested our hypothesis (Laverde et al., Chapter 3) that negative relationships expected under the transfer hypothesis are the result of sensory drive and need not involve tradeoffs in energy allocation; this hypothesis predicts that negative relationships between signals should be observed only when jointly considering birds from the understory and canopy due to the expected differences in communication strategies between habitats (Fig. 1e).

**Figure 1.**
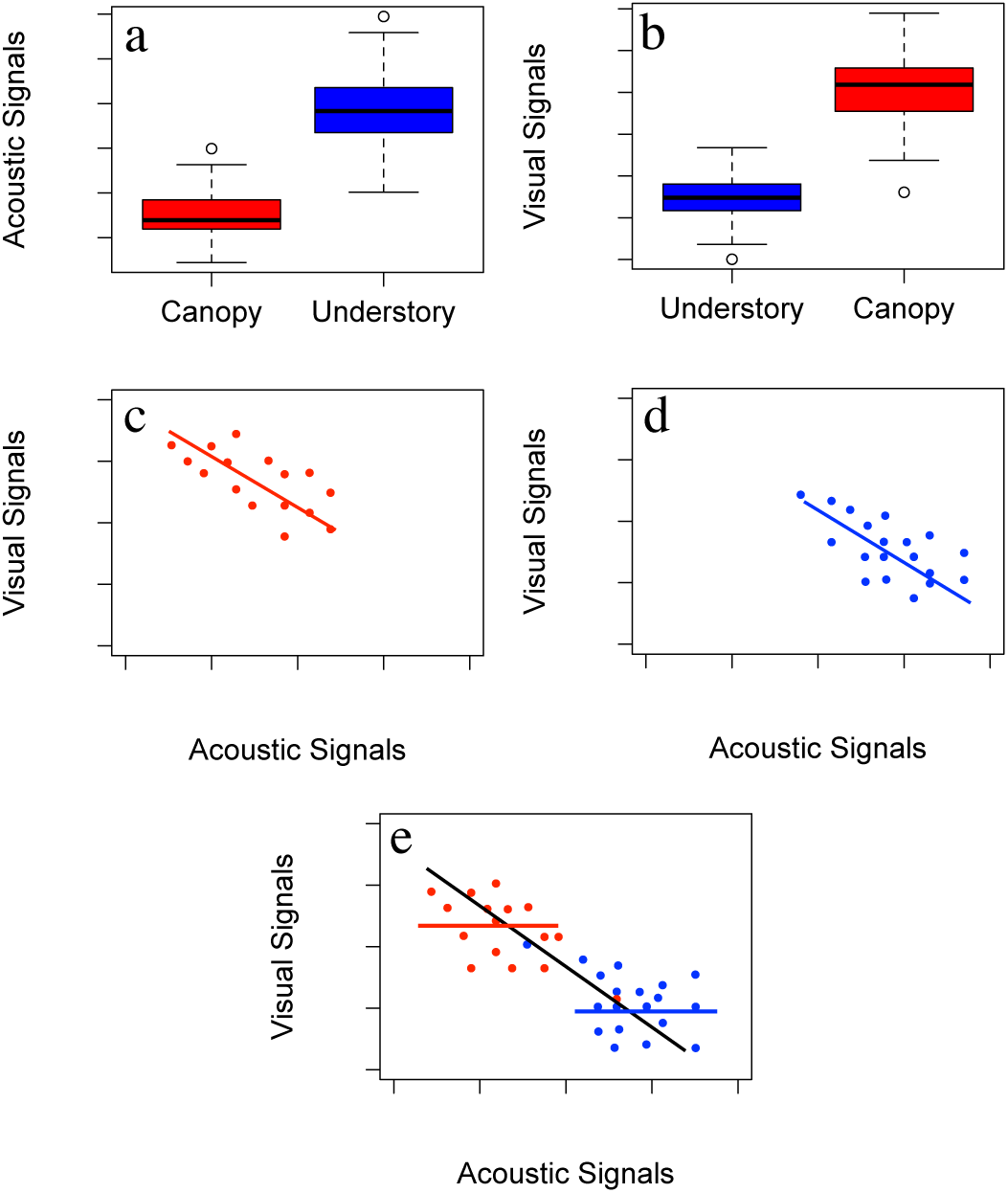
Predictions of the sensory drive and transfer hypotheses proposed to explain signal evolution. In our montane forest study site, the sensory drive hypothesis predicts that (a) visual signals should be most elaborate in the canopy where light availability is greater, and that (b) acoustic signals should be most elaborate in the understory where obstacles do not favor visual signals. The transfer hypothesis predicts (c and d) negative relationships between communication signals even in species from the same habitat. The hypothesis that sensory drive determines the negative relationships observed between communication signals predicts these relationships exist only when considering species from different habitats (e).

## Methods

We studied the bird assemblage in a tropical montane forest in Chingaza National Park, eastern Andes of Colombia. Chingaza is located ca. 40 km east of the city of Bogotá in the departments of Cundinamarca and Meta. The region has an average annual precipitation of 1800 mm, with two distinct peaks. The dry season extends from November to March with minimum rainfall in January and February, and the wet season extends from April to October with maximum rainfall in June and July (Vargas and Pedraza 2004). Approximately 190 bird species in 40 families occur in the park (Stiles and Rosselli 1998; Vargas and Pedraza 2004), but we focused on 52 species of 20 families that were common in forest in our study area (Table S1). We categorized each species as occurring in canopy or understory habitat based on our field observations and on information from the literature (Hilty and Brown 1986; Stiles and Rosselli 1998).

### QUANTIFYING INVESTMENT IN SONGS

We sought to characterize the investment of avian species in vocal signals by measuring (1) the frequency with which songs are emitted over relatively long time frames (i.e., vocal output), (2) the rate at which songs are delivered during singing (i.e., song rate), and (3) properties of individual songs, namely song length and syllable variety. Recordings for these measures were obtained by continuous sampling of avian vocalizations in six different locations in the Palacio sector of Chingaza National Park (4°41’ N, 73°50’ W; Table 1). Autonomous recording units (ARUs; Songmeter II, WildLife Acoustics) were located in forests between 2950 and 3170 m elevation and placed more than 500 m away from each other to avoid recording the same individuals in more than one unit. Each ARU was programmed to record for three minutes every 30 minutes. We sampled vocal output over several months in 2013: February (4 ARUs), April (3 ARUs), May (4 ARUs), July (2 ARUs), and August (2 ARUs; Table 1 and Table S1). We listened to recordings obtained from 5:30 am to 12:00 m (vocal activity decreased markedly later in the day) and identified vocalizing species (90% of avian vocalizations detected were identified to species).

**Table 1.** Phylogenetic analyses of variance of vocal variables with strata as a predictor. Statistical differences were found in variables related to vocal output as predicted by the sensory drive hypothesis (i.e. number of recordings counted in ARUs and number or recordings in sound collections/size of distributional range), but not in variables related to song elaboration. Significant differences after Bonferroni correction are shown in bold.

| Variable | F | p | N |
| --- | --- | --- | --- |
| Recordings in ARUs | 5.97 | **0.05** | 45 |
| Res. Sound collections/Area | 4.01 | 0.08 | 52 |
| Song Rate | 2.92 | 0.24 | 39 |
| Song Length | 3.16 | 0.24 | 37 |
| Syllable Variety | 0.21 | 0.64 | 37 |

We measured vocal output as the total number of recordings in which each species was present in the recordings obtained by the ARUs. Although data gathered by ARUs gave us information about the frequency with birds sang during our study period in Chingaza, they can not reveal patterns in vocal output over longer time frames. For example, monitoring during few months may be inadequate to characterize vocal output in species with markedly seasonal patterns of annual vocal activity (e.g. tinamous and thrushes). To obtain complementary information about vocal output over longer time frames, we also used data on recordings archived in sound collections; in a previous study we validated this approach by demonstrating that the number of recordings in archives reflects vocal output and temporal patterns in vocal activity measured through continuous monitoring (Laverde et al. Chapter 2). We thus tallied the total number of recordings of each of our study species archived in three major sound collections (Colección de Sonidos Animales at Instituto Alexander von Humboldt [CSA], Macaulay Library of Natural Sounds [MLS] and xeno-canto [XC]) corrected by the size of its distributional range (Laverde et al. Chapter 2) as an additional proxy of vocal output.

In addition to vocal output, we measured three variables related to song elaboration: song rate, song length, and syllable diversity (Cardoso and Hu 2011). We measured song rate as the mean number of songs of each species in the recordings in which we detected each species singing. We defined a song as a cluster of single vocal units separated from other clusters by a longer time span than that between any inter-unit time interval; songs may also be identified by the pauses between them, which may be on the order of several seconds (Thompson et al. 1994). We measured song length and counted the number of different types of syllables in Raven version 1.5 (Cornell Lab of Ornithology, Ithaca, New York, USA) using the following combination of settings: FFT window = 512, window type = Hann, and window overlap = 50%. We defined a syllable as the minimum discrete unit forming a song (Cardoso and Hu 2011); we recognized types of syllables by their shape on spectrograms and then counted them as our measure of syllable diversity. We measured syllable variety and song length using between 2 and 175 songs per species obtained from the xeno-canto archive (total 1352 songs; Table S2).

### QUANTIFYING PLUMAGE COLORATION AND CONSPICUOUSNESS

#### Colors of birds

To quantify plumage color heterogeneity and conspicuousness we measured 297 specimens of 52 species from 29 families (range 1-8 individuals per species; see suplementary material) in the ornithological collection of the Instituto de Ciencias Naturales, Universidad Nacional de Colombia. On each specimen we measured five color patches on different body parts (crown, rump, back, throat and belly) and also any other differently colored patch we observed in other body parts. Reflectance measurements were taken using an Ocean Optics USB-4000 Spectrometer and a DH-2000 deuterium halogen light source coupled with an optic fiber QP400-2-UV-VIS with a 400 um diameter. Light was reflected on the surfaces at a 45° angle. The spectrometer was calibrated using a white and a black standard before measuring each specimen. Each color patch was measured twice and measures averaged using the *pavo* package for R (Maia et al. 2013). We measured males and females for all species, but because none of the species was sexually dichromatic, we combined all the individuals for analyses.

We used the tetrahedral color space model (Endler and Mielke 2005; Stoddard and Prum 2008) to describe plumage coloration for each species. We used models of cone sensitivity spectra corresponding to the UV-sensitive (UVS) and V-sensitive (VS) types to account for the two main types of color vision in diurnal birds (Ödeen and Håstad 2009); because results were similar using both models, we only show results for UV-type vision. We considered two measurements indicating investment in visual signals as a basis for our tests of the transfer and sensory drive hypotheses: hue disparity and color conspicuousness. Hue disparity is an estimate of color heterogeneity, measured as the magnitude of the angle between two color vectors in the tetrahedral color space (Stoddard and Prum 2008). Color conspicuousness measures the mean contrast of plumage patches to the background (see below).

Determining how conspicuous avian plumages are requires characterizing the visual background in which species signal. Thus, we collected 35 leaves from different canopy trees along a 50 m transect, and randomly took 50 samples of litter and tree bark in the same transect in the understory to quantify background colors in the canopy and understory, respectively. We measured the reflectance spectra of these materials using the same equipment and protocol used to measure bird specimens. Each sample of each element (i.e. one leave, a piece of litter or bark) was measured twice and measurements were averaged. We then averaged measurements of all canopy leaves to obtain a mean canopy background reflectance spectrum, and averaged all the litter and bark measurements to obtain the mean background spectrum for the understory (Gomez & Thery, 2007). To quantify the chromatic contrast (ΔS) between plumage patches (crown, back, rump, throat and breast) and the background color we used the receptor noise model (Vorobyeb et al. 1998). This model considers the sensitivity of cones in the avian visual system and estimates of cone abundance and noise in photoreceptors to calculate discrimination thresholds or ‘just noticeable differences’ (JND). Values in the 1–3 range indicate that two objects are likely to be indistinguishable to an avian observer, whereas values >3 are increasingly likely to lead to detection and discrimination (Vorobyev et al. 1998). We contrasted each color patch in bird plumages against the color of leaves in the canopy or litter and tree bark in the understory depending on the habitat occupied by each species using the *coldist* function in *pavo*. Finally, we estimated overall color conspicuousness as the mean chromatic contrast of plumage patches against background color expressed in JND units.

### STATISTICAL ANALYSES

To evaluate the effect of habitat on the elaboration of visual and acoustic signals (i.e. sensory drive hypothesis) we tested the predictions that (1) colors are more diverse and more conspicuous in the canopy than in the understory, and that (2) vocal output is greater and songs are more elaborate in the understory than in the canopy. To evaluate these predictions we ran phylogenetic analyses of variance (ANOVAs) with habitat as predictor variable and measures of color and song elaboration as response variables. Owing to multiple comparisons we used the Bonferroni correction to adjust the p-values: this is a method in which the p-values are multiplied by the number of comparisons. To evaluate the transfer hypothesis we tested the prediction that measurements of elaboration in vocal and visual signals are negatively related using linear models and phylogenetic generalized least-square multiple regressions (PGLS), with hue disparity and color conspicuousness as response variables and song length, song rate, syllable variety and vocal output as explanatory variables. Finally, to evaluate the hypothesis that the negative relationships between songs and colors are the result of sensory drive operating separately on two types of communication signals we tested the prediction that negative relationships between signals should be observed only when considering birds from the understory and canopy together. We ran linear models and PGLS of understory and canopy birds separately, and including all species occurring in the understory and canopy. If negative relationships are due to the effect of sensory drive, then they should be observed only when both canopy and understory birds are included in analyses; negative relationships between signals in species from the same forest stratum would be consistent with tradeoffs. All PGLS analyses were ran using the caper package for R (Orme et al. 2012) based on a comprehensive avian phylogeny (Jetz *et al.* 2012). Because we obtained color measurements for our 52 study species but were able to obtain good quality recordings only for 38 species for which we quantified song length and syllable variety, our final analyses of the transfer hypothesis focused on 38 species having complete data for colors and songs (Table S2).

Some of our results depended on whether or not analyses accounted for phylogenetic effects (see below). Therefore, we explored variation in vocal output and measurements of elaboration of visual signals (i.e. hue disparity and color conspicuousness) between understory and canopy birds separately for species in different taxonomic groups (non-passerines, and suboscines and oscine passerines). For these analyses, we ran linear models setting forest stratum and taxonomic group as predictors.

## Results

As predicted by the sensory drive hypothesis, vocal output (measured by our continuous field monitoring using ARUs) differed between forest strata: understory birds sing more frequently than canopy birds (Table 1, Fig. 2), but the number of recordings in sound collections were marginally significant (Table 1, Fig. 2). By contrast, variables related to song elaboration (i.e. song rate, song length and syllable variety) did not differ significantly between understory and canopy species (Table 1, Fig. 2).

**Figure 2.**
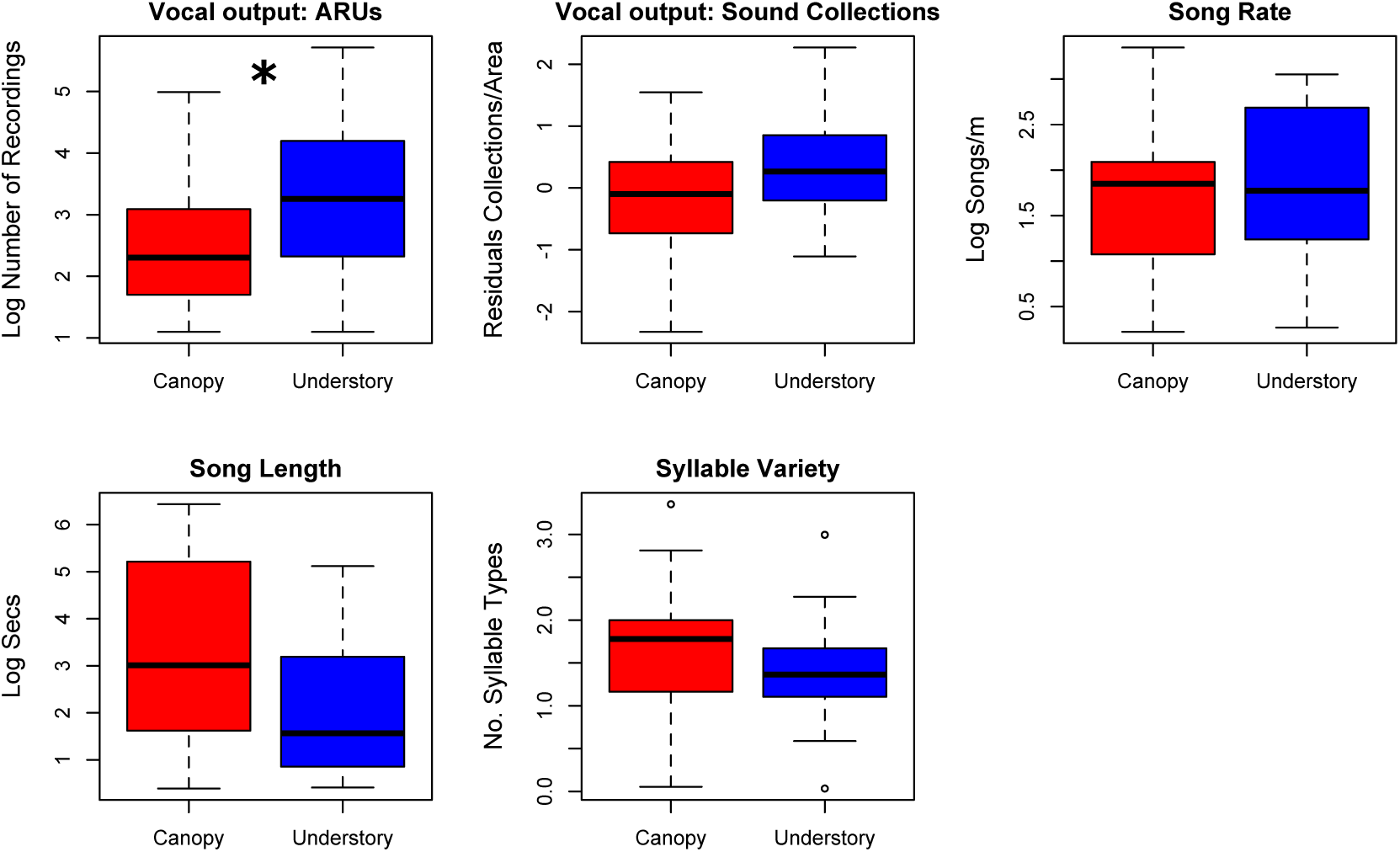
Variation in acoustic communication strategies between understory and canopy bird species from Chingaza National Park (see sample sizes in Table 1). Boxplots show two measures of vocal output (recordings counted in autonomous recording units ARUs and in three sound collections) and four measures of song elaboration: song rate, song length, syllable variety and syllable diversity. Analyses indicate that understory birds sing more frequently than canopy birds in the recordings collected in ARUs as predicted by sensory drive, but the numbers in sound collections were not statistically significant. There were not differences between habitats in any measure of song elaboration. Significance of statistical results of the phylogenetic ANOVAs after Bonferroni correction: * p<0.05.

Variation in visual signals partially supported predictions of the sensory drive hypothesis. Although plumage hue disparity did not differ between understory and canopy birds, the contrast of two color patches (i.e. back and breast) to the background and color conspicuousness were greater in canopy than in understory species as we had predicted (Table 2, Fig. 3). The contrast of crown, throat and rump color with the background did not differ between understory and canopy species.

**Table 2.** Phylogenetic analyses of variances for the color variables with habitat as a categorical predictor. Statistical differences were found in the contrast to background of two patches (i.e. back and breast) and color conspicuousness as predicted by the sensory drive hypothesis. The color of throat, rump, crown and hue disparity were not statistically different between understory and canopy birds. Significant differences after Bonferroni correction are shown in bold.

| Variable | F | p | N |
| --- | --- | --- | --- |
| Crown contrast to BG | 3.58 | 0.12 | 52 |
| Back contrast to BG | 5.52 | **0.04** | 52 |
| Rump contrast to BG | 3.87 | 0.15 | 52 |
| Breast contrast to BG | 5.22 | **0.04** | 52 |
| Throat contrast to BG | 0.03 | 0.96 | 52 |
| Color Conspicuousness | 4.624 | **0.05** | 52 |
| Hue Disparity | 0.67 | 0.82 | 52 |

**Figure 3.**
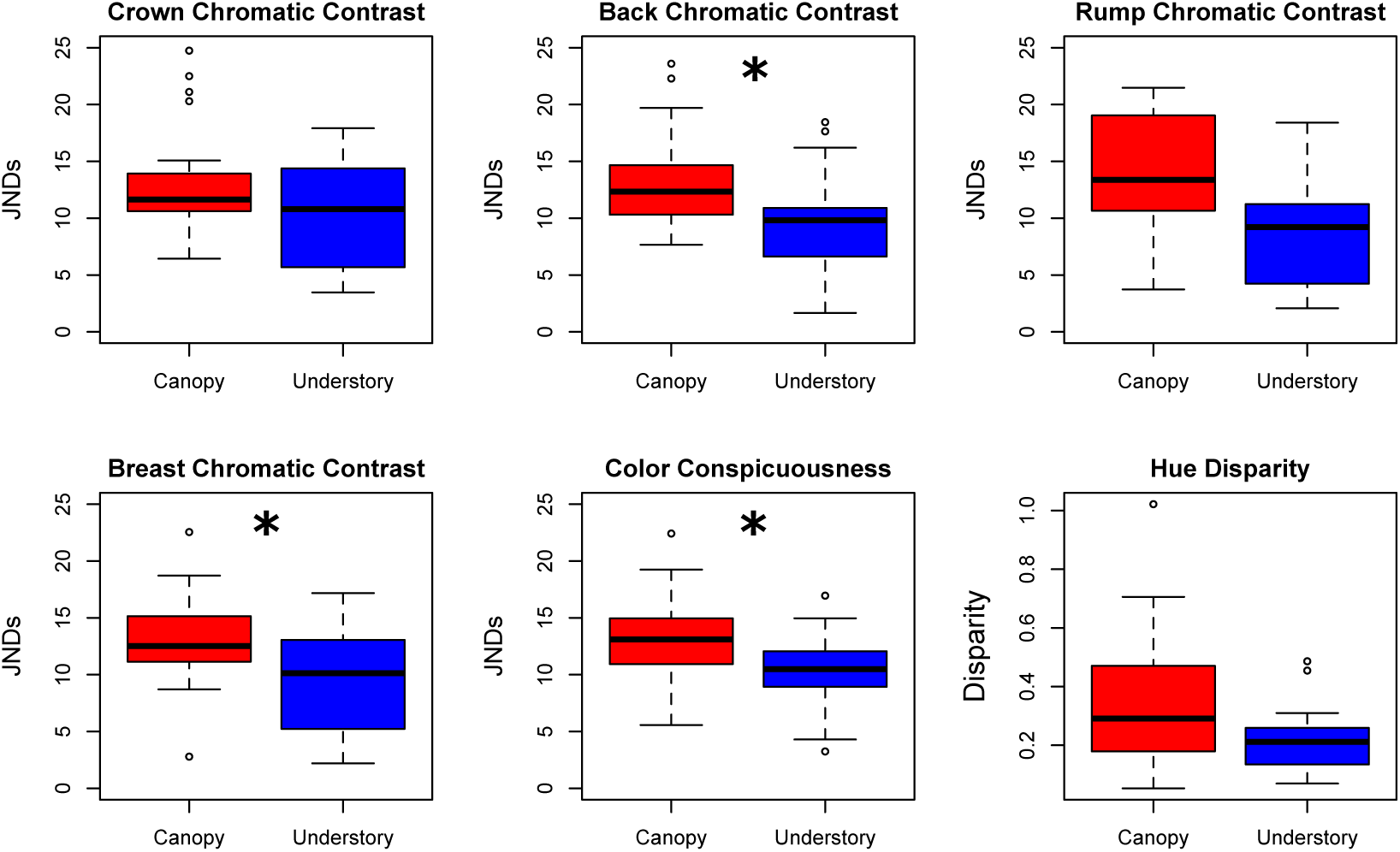
Variation in visual communication strategies between understory and canopy bird species from Chingaza National Park (see sample sizes in Table 2). Boxplots show measures of the contrasts of patches to background, color conspicuousness and hue disparity. Analyses indicate that understory canopy birds contrast more to the background than understory birds, although the crown, throat and rump were not different after Bonferroni correction. Hue disparity was not different between strata, but variation was greater in the canopy. Chromatic contrasts were measured as just noticeable differences (JNDs, see text). Significance of statistical results of the phylogenetic ANOVAs: * p<0.05.

Phylogenetic multiple regressions including our two measures of vocal output (number of recordings obtained by continuous monitoring using ARUs and the total number of recordings found in three sound collections), song length, syllable variety, and song rate as predictors of plumage hue diversity and conspicuousness revealed significantly negative relationships irrespective of habitat consistent with the transfer hypothesis. However, the most relevant variables explaining variation in our measures of investment in color signals differed between models (Table 3 and 4). Color conspicuousness was negatively related to the number of recordings in ARUs (Table 3): birds with colors contrasting less against the background sing more frequently. In turn, hue disparity was negatively related to song length (birds with lower diversity of colors produce longer songs) but not to any other measure of vocal output or song elaboration (Table 4).

**Table 3.** Phylogenetic generalized least squares multiple regressions of color conspicuousness on a set of predictors related to vocal output and song elaboration (F = 3.34, R^2^ = 0.25, p = 0.01. AIC=24.03, N=28). Color conspicuousness was negatively related to the number of recordings in ARUs: birds with colors contrasting less against the background sing more frequently, as predicted by the transfer hypothesis. Significant difference is shown in bold.

| | $\beta$ | t-value | p |
| --- | --- | --- | --- |
| **ARUs** | **-1.32** | **-2.55** | **0.01** |
| Syllable Variety | 0.79 | 0.97 | 0.34 |
| Song Length | 0.52 | 1.24 | 0.22 |
| Song Rate | 0.54 | 1.35 | 0.18 |

**Table 4.** Phylogenetic generalized least squares multiple regressions of hue disparity on a set of predictors related to vocal output and song elaboration (F = 3.32, R^2^ = 0.24, p = 0.02, AIC = 46.98, N=28). Song length was negatively related to hue disparity: birds with lower diversity of colors produce longer songs as predicted by the transfer hypothesis. Significant difference is shown in bold.

|  | **$\beta$** | **t-value** | **p** |
| --- | --- | --- | --- |
| ARUs | -0.15 | -1.78 | 0.08 |
| Syllable Variety | 0.05 | 0.41 | 0.68 |
| **Song Length** | **-0.17** | **-2.46** | **0.02** |
| Song Rate | 0.05 | 0.83 | 0.41 |

As predicted by our hypothesis that negative relationships as those expected under the transfer hypothesis are the result of sensory drive (Fig. 4), linear regressions between hue disparity and our two measures of vocal output were significantly negative only when we considered understory and canopy birds together (Table 5); when we ran separate analyses for understory and canopy birds we did not find any relationship.

**Table 5.** Lineal regressions and PGLS of hue disparity and color conspicuousness and our two measures of vocal output. Results of linear models with hue disparity as response variable supported our hypothesis that negative relationships between signals are observed when considering birds from the understory and canopy due to the expected differences in communication strategies between habitats. PGLS models partially supported this hypothesis, but also the negative and significant relationships found within the understory supported the transfer hypothesis. All the significant relationships marked with an asterisk were negative.

|  | Lineal Regressions |  |  | PGLS models |  |  |
| --- | --- | --- | --- | --- | --- | --- |
|  | All<br>Birds | Canopy<br>Birds | Understory<br>Birds | All<br>Birds | Canopy<br>Birds | Understory<br>Birds |
| Hue Disp. ~ Rec. in ARUs | * | NS | NS | ** | NS | * |
| Hue Disp. ~ Rec. In Collections | * | NS | NS | NS | NS | * |
| Color Consp. ~ Rec. In ARUs | NS | NS | NS | NS | NS | NS |
| Color Consp. ~ Rec. In Collections | NS | NS | NS | NS | NS | NS |

\* p<0.05, \*\*p<0.01

**Figure 4.**
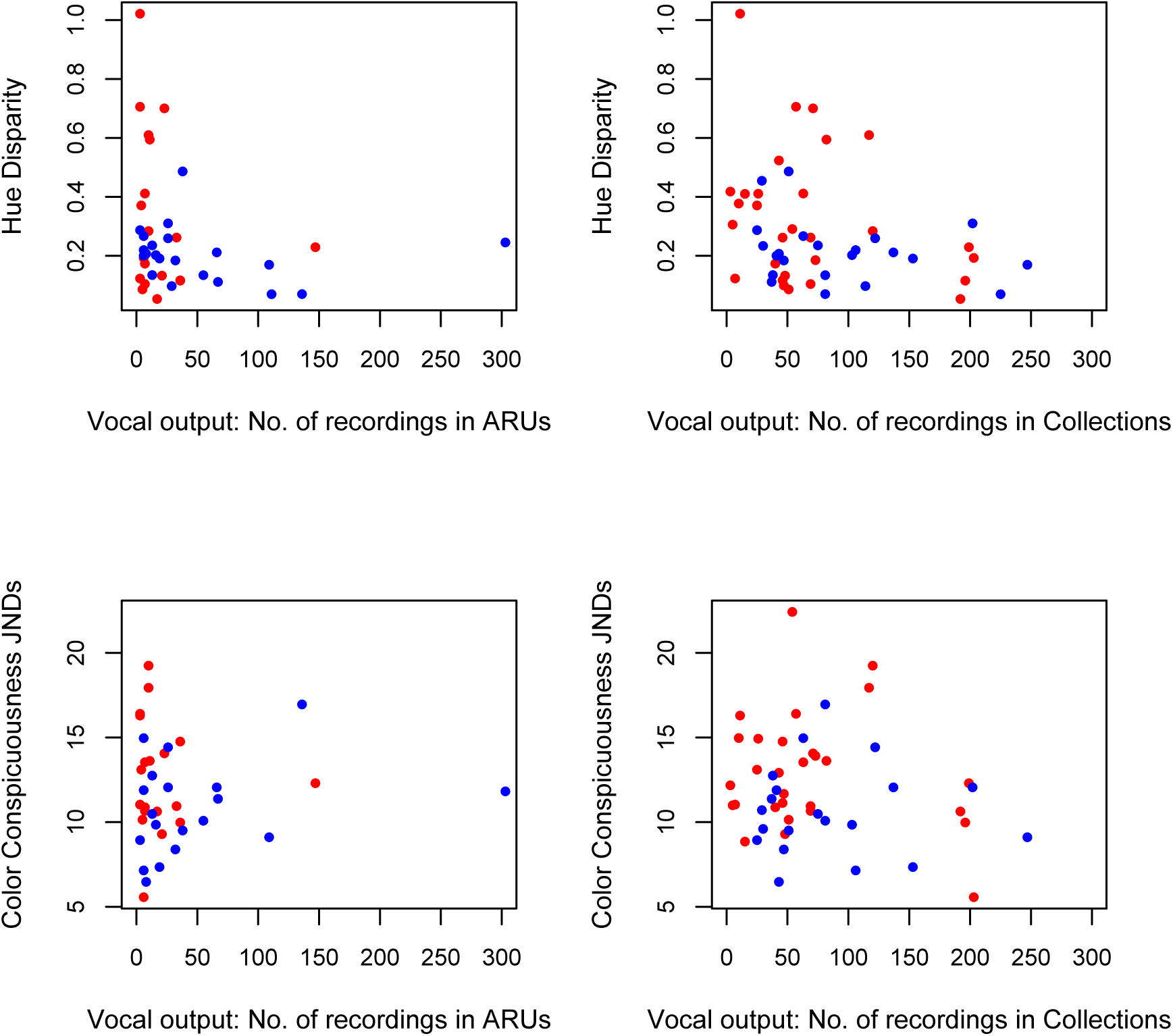
Relation between two variables describing investment in plumage coloration (i.e. hue disparity and color conspicuousness) and vocal output (i.e. number of recordings in ARUS and number of recordings found in sound collections). Understory species are shown in blue and canopy species in red. Notice that understory birds exhibit more variation and reach higher values in vocal output, whereas canopy birds have more variation and higher values in the two color variables studied.

As predicted by our hypothesis that negative relationships as those expected under the transfer hypothesis are the result of sensory drive (Fig. 4), linear regressions between hue disparity and our two measures of vocal output were significantly negative only when we considered understory and canopy birds together (Table 5); when we ran separate analyses for understory and canopy birds we did not find any relationship. However, PGLS gave slightly different results: we found negative significant relationships only between hue disparity and the recordings obtained by the ARUs (Table 5), but not between hue disparity and the number of recordings archived in sound collections. As expected given our hypothesis that negative relationships in the investment in different types of signals are mediated by habitat, hue disparity and our measures of vocal output were unrelated in canopy birds; however, in understory birds these variables were negatively related, a pattern consistent with tradeoffs (Table 5). We found no significant relationships between measures of vocal output and color conspicuousness when considering canopy and understory birds together, neither when we ran separate analyses for canopy and understory birds (Table 5). Finally, when examining relationships between vocal output and visual signals separately for major taxonomic groups, we found different patterns in oscines and suboscines (Fig. 5). In oscines, vocal output (number of recordings in ARUs, t=-2.54, p=0.02; number of recordings in collections, t=-2.69, p=0.01) and color conspicuousness (t=3.29, p=0.004) were significantly different between strata in the direction we had predicted (i.e. greater vocal output in the understory and more conspicuous plumages in the canopy), but this was not the case for suboscines (number of recordings in ARUs, t=-0.90, p=0.38; number of recordings in collections, t=0.09, p=0.92; Hue disparity, t=-0.89, p=0.38, Conspicuousness, t=-0.69, p=0.49). Data for non-passerines were too limited to draw any definitive conclusions.

**Figure 5.**
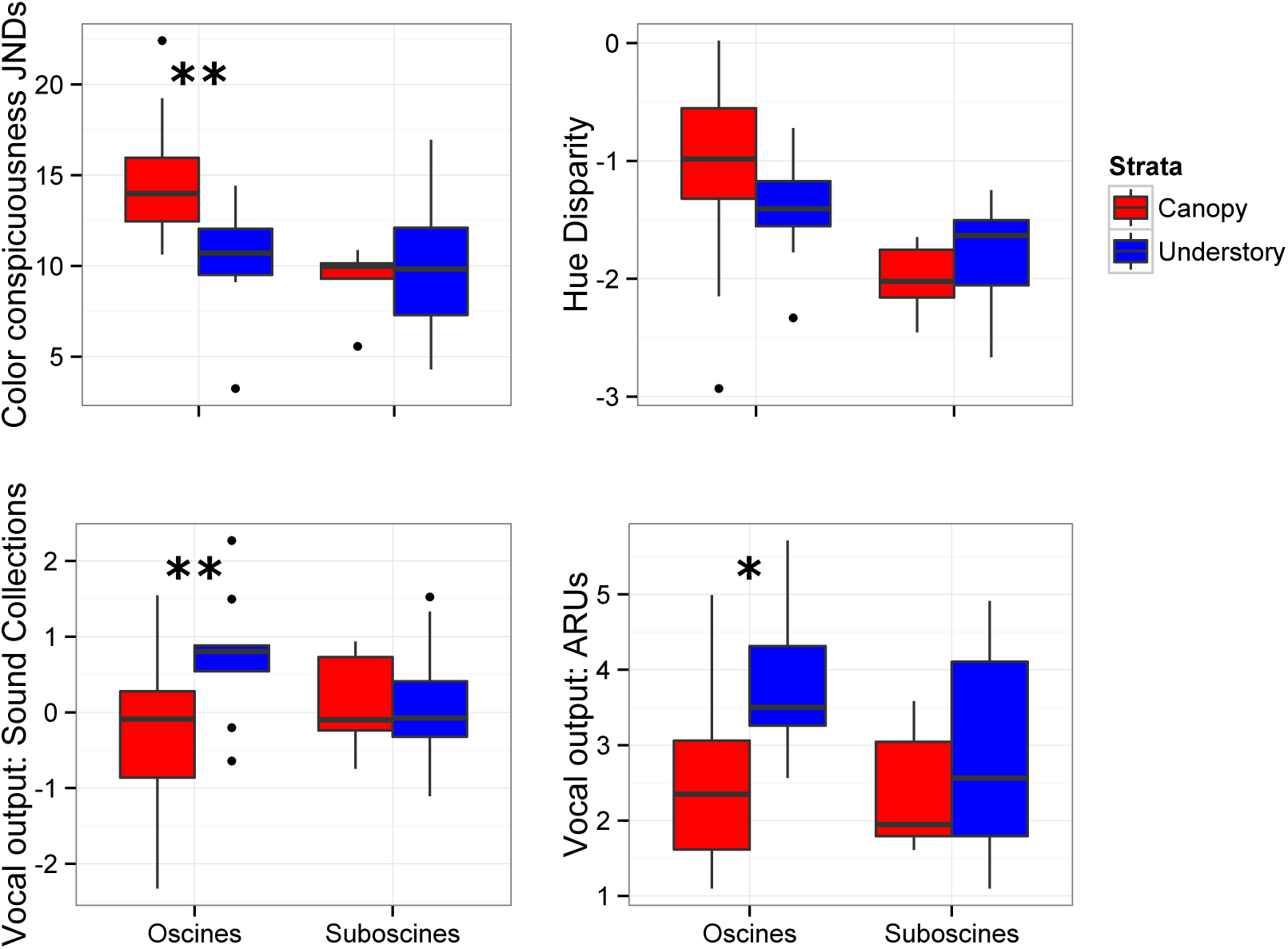
Boxplots showing variation in vocal output (measured using ARUs and the number of recordings in sound collections corrected by area of distributional ranges), hue disparity and color conspicuousness in understory and canopy birds belonging two taxonomic groups. In oscines (n=27 species) investment in acoustic and visual signals differs between habitats as predicted by sensory drive (birds are less conspicuous and sing more often in the understory than in the canopy), but this pattern was not present in suboscines. (n=17 species). Due to small sample sizes, non-passerines we excluded from this analysis. Statistical results of a t-student test * p<0.05, ** p<0.001.

## Discussion

Our work demonstrates that in a tropical montane forest assemblage, birds from the understory sing more frequently, whereas birds from the canopy exhibit more diverse and more conspicuous colors in their plumage. These two different signaling strategies are consistent with predictions of the sensory drive hypothesis and likely represent adaptations to maximize the perception of signals in two contrasting habitats (Endler 1992; Seehausen et al. 2008; Tobias et al. 2010). Our work goes a step further relative to other tests of the sensory drive hypothesis by linking it to another leading explanation for signal evolution, namely the transfer hypothesis (Gilliard 1956; Shutler and Weatherhead 1990; Shutler 2011). Our study (see also Laverde et al. Chapter 3) suggests that negative relationships between songs and colors observed by early naturalists that led to the postulation of the transfer hypothesis (Darwin 1871; Huxley 1938; Gilliard 1956) do not necessarily arise as a result of tradeoffs. Rather, such negative relationships may be a consequence of selection for optimal signaling strategies in the local habitat of each species.

The sensory drive hypothesis postulates that natural selection favors signals, receptors, and signaling behavior that maximize signal reception relative to background noise and minimize signal degradation (Endler 1992). Remarkably, most of the studies on this topic have evaluated changes in signal design and in sensory systems in relation to properties of habitats (e.g., Seddon 2005; Seehausen et al. 2008; Tobias et al. 2010), but we are aware of a single study examining the potential consequences of sensory drive on behavioral traits associated to signaling (i.e. courtship behavior in sticklebacks, Heuschele et al. 2009). Our work shows that not considering aspects of signaling behavior may lead to incorrect inferences about the validity of the sensory drive hypothesis. Specifically, had we not considered behavior we would have likely rejected the sensory drive hypothesis because, contrary to its predictions, variables associated to the elaboration of songs (song length and syllable variety) were not affected by habitat in our study system. However, we found differences in vocal output between understory and canopy birds in the direction predicted by sensory drive: understory birds sing significantly more often than canopy birds, likely because producing vocalizations more frequently may increase the chance of the reception of acoustic signals in the dense understory of montane tropical forests. Because vocal output can be measured relatively easily using ARUs and can be reasonably approximated using data in sound collections (Laverde et al. Chapter 2), we suggest that future studies testing for sensory drive should consider such measures as an important dimension along which animal signaling may respond adaptively to environmental variation. For example, in other avian groups like trogons (Ornelas et al. 2009) and tanagers (Mason et al. 2014) there is seemingly no negative relation between investment in song and in colors, but these studies only focused on variables associated to elaboration of songs and did not consider variables related to vocal output. Examining vocal output as an indication of investment in vocal signals may reveal negative relationships between types of signals in these groups.

The features related to song elaboration (i.e. song length, syllable diversity and song rate) were not significantly different between canopy and understory birds. Songs of birds contain multiple traits that can be sexually selected, which are not equally important for all species: females of some species prefer males that produce very diverse repertoires, but females of other species may prefer longer songs or higher song rates (Gil and Gahr 2002). Even within the same group of birds (i.e. wood warblers), females can select different traits of songs, sometimes resulting in simple signals (Cardoso and Hu 2011). Therefore, an assemblage-base approach may place together a diversity of preferences from different species. The classical hypotheses proposed to explain the evolution of complex songs assumed to be the outcome of sexual selection acting on males (Price 2015), but in the tropics female bird song is much more common than previously thought, suggesting that the classical view of sexual selection acting on the complexity or elaboration of male songs, must be re-evaluated (Odom et al. 2014; Price 2015).

We also found evidence consistent with sensory drive in plumage coloration; overall, canopy birds contrasted more relative to the background color than understory birds, a result consistent with previous work on a lowland assemblage (Gomez & Thery 2007). In addition, although the diversity of colors (i.e. mean hue disparity) was not different between forest strata, variation in color among canopy birds (coefficient of variation [cv]=69.5) was wider than variation among understory birds (cv=47.8). In the understory, the majority of the birds are mostly cryptic and not very colorful, but in the canopy one can find species with dull and uniform plumages (e.g. Smoky Bush-tyrant *Myiotheretes fumigatus,* Great Thrush *Turdus fuscater*) or conspicuous/colorful (many tanagers) plumages. We suggest that exhibiting colorful plumages in the canopy might not be as risky as in the understory due to varying predation pressure between strata (Gotmark and Post 1996). However, more data on abundance of predators, their distribution and their effects on the evolution of communication signals is required to effectively link variation in signals with habitat-specific predation pressures, especially in the tropics.

In principle, our results support the idea of transference between communication signals: birds less contrasting against the background sing more often, and birds with more diversity of colors produce shorter songs. Furthermore, the negative relationship we observed between vocal and visual signals of understory birds seems to support the idea of classical tradeoffs: hue disparity and our two measures of vocal output were negatively related when accounting for phylogeny. Nevertheless, we argue that the effect of habitat is without doubt a crucial factor shaping communication signals and some of the observed negative relationships between them.

Suboscine and oscine passserines showed different responses to the effect of the habitat with regard to visual and acoustic signals. In oscines, songs and colors varied between habitats as predicted by sensory drive (songs more important in the understory, colors in the canopy), but this was not the case in suboscines. Suboscines and oscines exhibit striking differences in habitat, diet, behavior, syrinx complexity and song development (Kroodsma 1984; Ricklefs 2002) which may account for the different patterns in signal evolution documented by our study. Foremost, vocal learning is crucial to the production of acoustic signals in oscines, but it is thought to be absent in the majority of suboscines, in which songs are developed without an influence of experience (Kroodsma 1984, 2007; Touchton et al. 2014). In addition, the greater complexity of the oscine syrinx allows for greater versatility (hence presumably greater evolutionary lability; see Weir and Wheatcroft 2011) in sound production relative to suboscines (Kroodsma 2007; Amador et al. 2008). Ecology may also help to explain the different patterns observed in oscines and suboscines. In the Neotropics, oscines dominate the luminous forest canopy and habitats outside the forest, but most of the birds of the dark forest understory belong to the suboscines, although a few suboscines (i.e. particularly flycatchers) have colonized the canopy and open areas (Ricklefs 2002). Most suboscines feed largely on insects whereas oscines make greater use of fruit resources (consistent with their dominance in the canopies of forests; Ricklefs 2002) and this may reflect on the ease with which plumages may evolve adaptively in response to habitat variation because various elements of visual displays are based on carotenoid pigments obtained form the diet and deposited on feathers (Hill and McGraw 2006). In sum, we hypothesize that vocal learning and greater complexity in anatomical structures involved in producing vocalizations coupled with a more diverse color palette associated with diets rich in carotenoids jointly allow for greater opportunities for adaptive evolution in oscine signals. This may partly account for our results showing signal evolution is consistent with sensory drive in oscines but not suboscines. However, other studies on suboscines have provided evidence that sensory drive shapes spectral and temporal properties of songs in this clade (Seddon 2005, Tobias et al. 2010).

To conclude, we suggest that the study of the evolution of animal communication signals needs to integrate traits indicative of signaling behavior (i.e. vocal output) to the classical view of only studying signal elaboration or complexity. In our study, understory birds emit more vocalizations than canopy birds, implying that habitat drives changes in a behavioral trait associated to signaling although signal elaboration *per se* may not differ between environments. More generally, considering that communication is multimodal, that signals do not act independently, and that many animals communicate by the integration of signals from different sensory modalities (Hebets and Papaj 2004), it is remarkable that much past research on the evolution of animal communication systems has largely focused on single and isolated signals. An overarching conclusion of our study is that examining multiple communication channels and considering their efficiency in local environments is essential to understand the mechanisms underlying the evolution of communication signals.

## Acknowledgments

We thank Chingaza National Park for allowing us to record bird songs. Greg Budney and Matt Medler of the Macaulay Library of Natural Sounds at Cornell University, and Paula Caycedo of the Colección de Sonidos Ambientales of the Instituto Alexander von Humboldt gave us unrestricted access to the information archived in their collections. We thank all the recordists who have deposited their recordings in these archives and who have made them available through xeno-canto. We are very grateful to Laura Céspedes who made all the field measurements of the background color. We are grateful to Simón Quintero, Ignacio Areta, Mecky Hollzmann, Trevor Price and Alex Kirschel for asistance in the field. This work was supported by funds from the Facultad de Ciencias at Universidad de los Andes, Bogotá, Colombia and Fundación Biodiversa-Colombia.

---

**Table S1.** Number of recordings of avian vocalizations gathered using ARUs by month and elevation in Chingaza National Park from February to August 2013.

| **ARU**<br>**No** | **February** | **April** | **May** | **July** | **August** | **Total** | **Geographical**<br>**Coordinates** | **Elevation (m)** |
| --- | --- | --- | --- | --- | --- | --- | --- | --- |
| **1** | 248 |  |  |  |  | 248 | 4°41'47"; 73°50'53" | 2950 |
| **2** | 121 | 138 | 92 |  |  | 351 | 4°41'54"; 73°50'50" | 2960 |
| **3** | 158 |  |  |  |  | 158 | 4°42'08"; 73°50'50" | 3015 |
| **4** | 233 |  | 121 | 246 | 246 | 846 | 4°41'44"; 73°50'39" | 3030 |
| **5** |  | 88 | 38 |  |  | 126 | 4°41'27"; 73°50'24" | 3015 |
| **6** |  | 131 | 132 | 150 | 50 | 463 | 4°42'42"; 73°50'21" | 3170 |
| **Total** | 760 | 357 | 383 | 396 | 296 | 2192 |  |  |

**Tabla S2.**
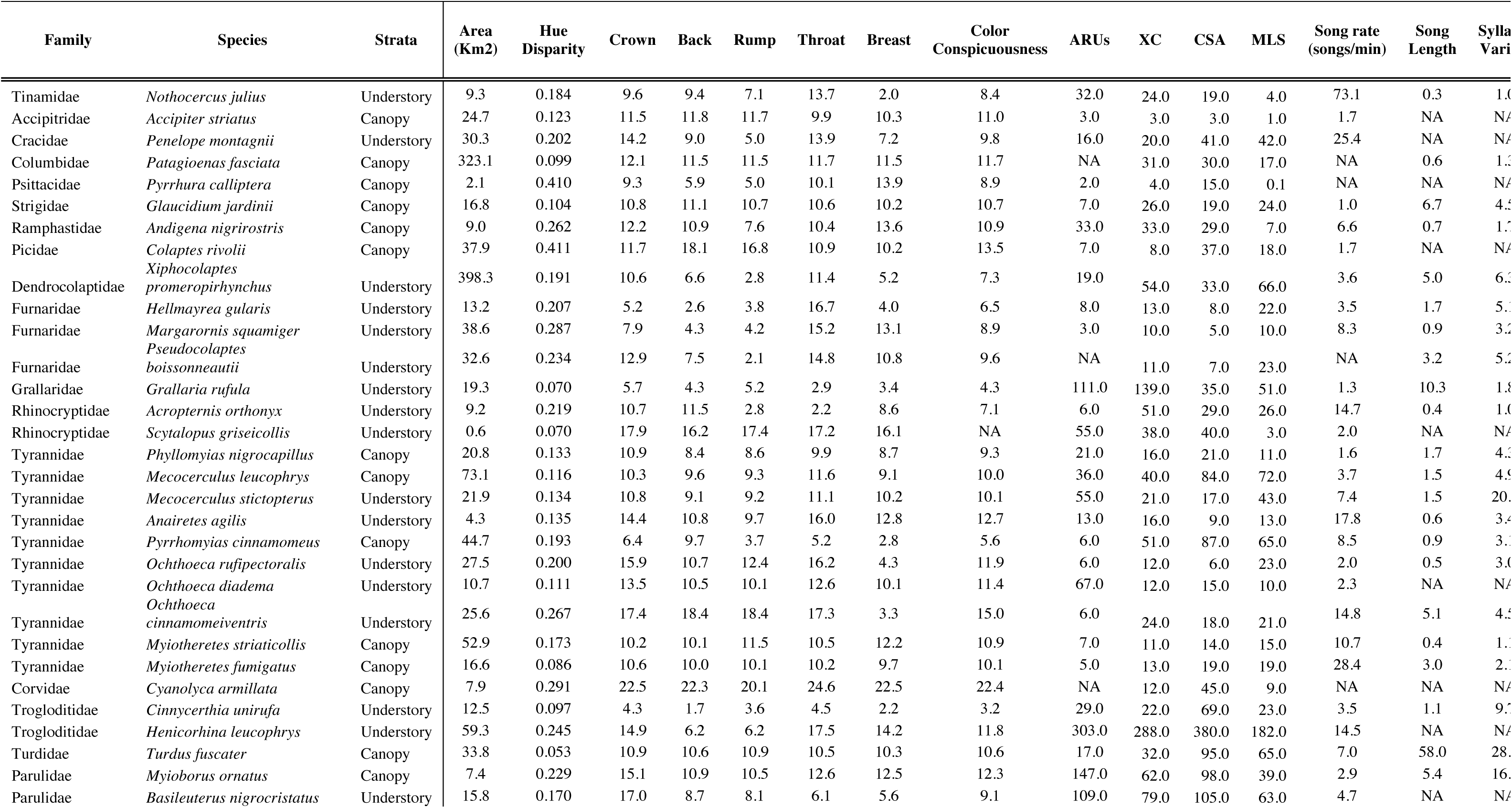
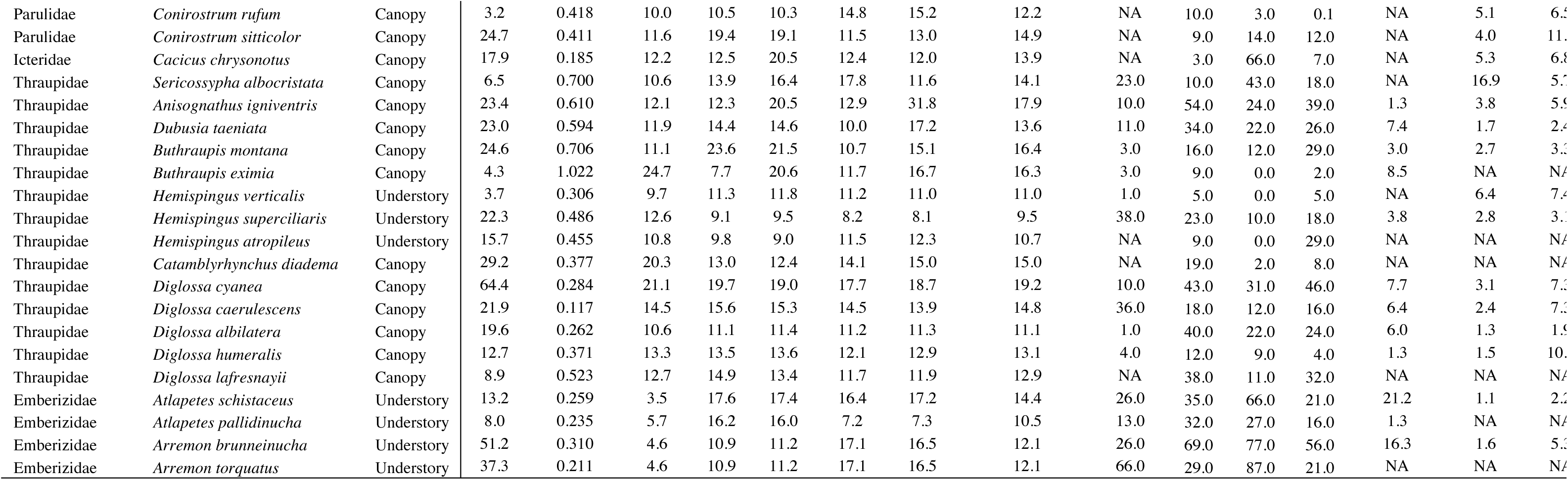
Data on habitat, area of distribution, hue disparity, color conspicuousness and vocal elaboration and output for 52 species present in the Palacio sector in Chingaza National Park.

